# gapsplit: Efficient random sampling for non-convex constraint-based models

**DOI:** 10.1101/652917

**Authors:** Thomas C. Keaty, Paul A. Jensen

## Abstract

**Summary:** Gapsplit generates random samples from convex and non-convex constraint-based models. Gapsplit targets under-sampled regions of the solution space for uniform coverage.

**Availability and Implementation:** Python and Matlab source code are freely available at http://jensenlab.net/tools.

## Introduction

Constraint-based models allow systems-level interrogation of biochemical networks with minimal kinetic data. The number of variables in constraint-based models far exceeds the number of kinetic parameters, resulting in underdetermined systems of equations that produce an infinite number of solutions. Rather than focus on any single solution, modelers can use ensembles of randomly sampled solutions to analyze network properties (Schellenberger and Palsson, 2009).

The leading algorithms for sampling constraint-based models use the “hit-and-run” (HR) framework (Smith, 1984). HR sampling walks through a model’s solution space by randomly selecting directions based on a set of warmup points. HR’s efficiency is tied to the convexity of the solution space. Since a convex combination of any number of existing solutions is also a solution, HR algorithms can quickly generate new solutions without resolving the model. When applied to constraint-based models, current HR samplers (ACHR (Kaufman and Smith, 1998), OptGP (Megchelenbrink et al., 2014), and CHRR (Haraldsdóttir et al., 2017)) quickly generate a series of samples that converge to a stable distribution. However, models containing reactions with fixed bounds can drastically reduce the fraction of the total sample space covered by the HR samplers (see Binns et al. (2015) and data below). One random sampler with improved coverage uses a “poling” method to push the random walk of the HR sampler away from previous samples (Binns et al., 2015). While the poling method improved coverage, the resulting optimization problems are nonlinear and require orders of magnitude more computation time.

Adding transcriptional regulation or enzymatic complexes to a model requires discrete variables, making the model non-convex. HR samplers cannot directly sample non-convex models. As a workaround, the ll-ACHR sampler uses a boxing approximation to sample models with (non-convex) loopless flux constraints (Saa and Nielsen, 2016). The box constraints enclose the non-convex solution space with a convex hull, and the convex hull, not the original solution space, is sampled. Care must be taken to reject any infeasible samples that lie outside the solution space but inside the convex hull. The efficiency of boxed models also decreases with additional discrete variables or non-convex constraints (Kiatsupaibul et al., 2011).

We present a new class of random sampler for constraint-based models. Our algorithm – called Gapsplit – uses mathematical programming to find solutions in the underexplored areas of the model’s solution space. Unlike HR algorithms, Gapsplit samples convex and non-convex models directly. Samples identified by Gapsplit uniformly cover a model’s solution space. Gapsplit yields better coverage than HR samplers for tightly constrained and non-convex models.

## The Gapsplit sampler

Gapsplit is designed to find sample points that uniformly cover the entire solution space. Gapsplit’s objective is to minimize the size of each variable’s *max gap*, the largest interval between two adjacent sample points (Figure 1A). Given a set of samples, Gapsplit selects a single variable and identifies its max gap. Gapsplit adds a constraint requiring the next solution be in the center of the max gap (the *target*; see Figure 1A). The model is solved to find such a solution, the constraint is removed, and the process is repeated with a different variable. To speed up sampling, Gapsplit also attempts to simultaneously split the max gaps of *k* randomly selected variables. Gapsplit uses a quadratic objective function to minimize the distance between the next solution and the centers of the max gaps for the *k* other reactions. (A complete description of the Gapsplit algorithm is presented in the Supplementary Methods.) The Gapsplit algorithm can be applied to any mathematical program including models with binary or integer constraints.

**Figure 1.**
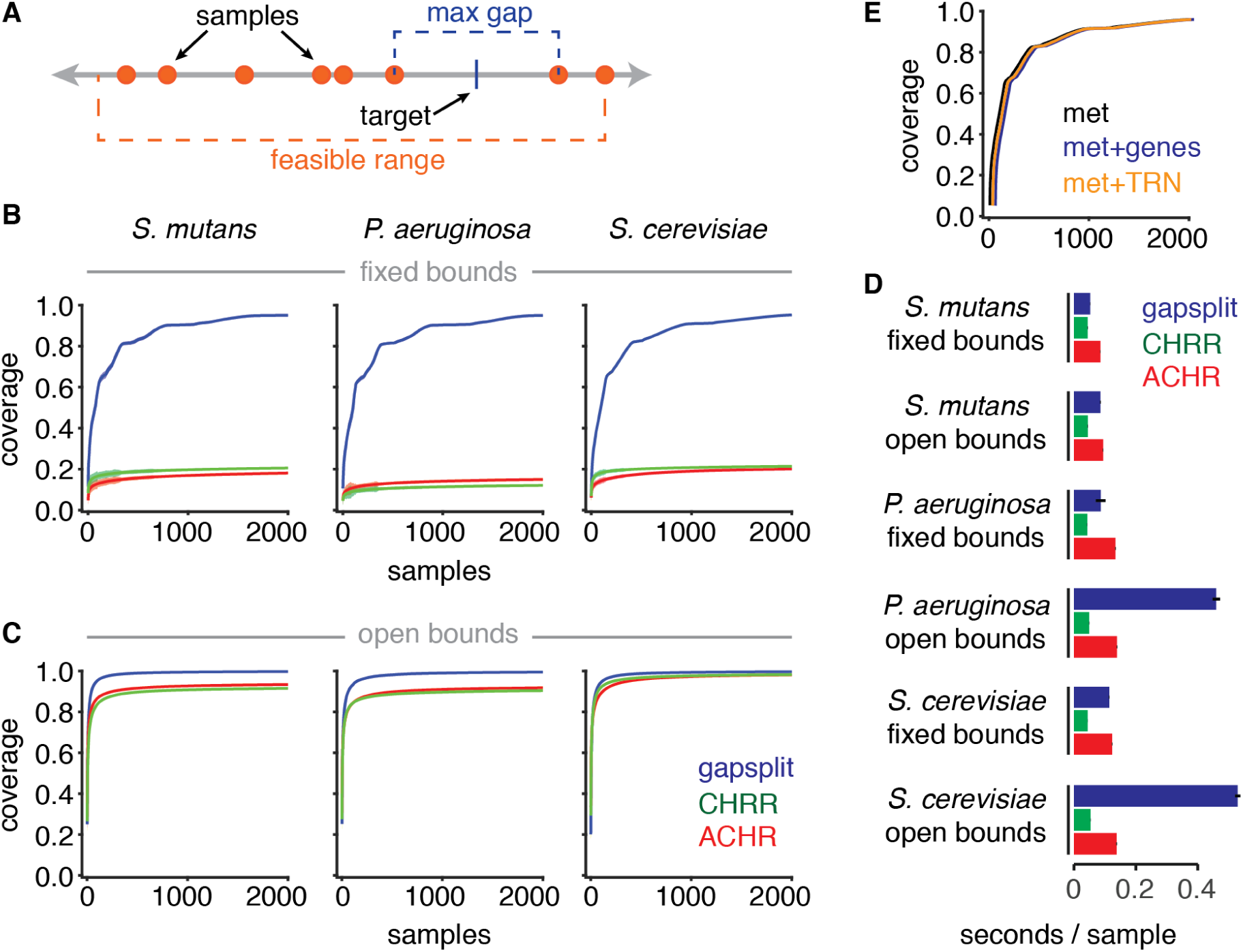
**A.** Gapsplit finds random samples for each variable (orange circles) within the variable’s feasible range. If a variable is selected for targeting, the next solution will be at a target that splits the maximum remaining gap in half. **B.** Gapsplit yields better coverage with fewer samples than HR family samplers. The mean (solid line) and 95% confidence intervals (color shading) are shown for 100 runs of each sampler on three metabolic models: bacteria *Streptococcus mutans* (iSMU v1.0, Jijakli and Jensen (2018)) and *Pseudomonas aeruginosa* (iMO1056, Oberhardt et al. (2008)); and the yeast *Saccharomyces cerevisiae* (iND750, Duarte et al. (2004)). Bounds were fixed to glucose minimal media as specified in the original publications. **C.** The coverage of HR family samplers improves if the bounds on exchange reactions are relaxed to arbitrarily large values. However, such conditions are not physiologically reasonable. **D.** Gapsplit is slower than the CHRR but faster than the ACHR samplers when models have fixed bounds. Mean time per 1000 samples is shown for 25 independent runs of each algorithm. Error bars show the standard deviation. Opening the bounds on the model slows all three algorithms. **E.** Gapsplit can sample non-convex models. Adding binary constraints for gene associations (met+genes, blue) or logical constraints for transcriptional regulation (met+TRN, orange) does not affect sampling of the yeast metabolic model (met, black). Each line represents the mean coverage for 50 independent simulations.

## Results

One metric to assess the quality of a set of random samples is coverage, defined as

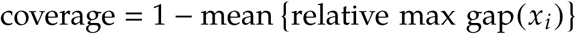

where *x*_*i*_ is a variable in the model and the *relative max gap* is the max gap of *x*_*i*_ divided by the feasible range of *x*_*i*_. Coverage ranges from 0 (all points at the edges of the feasible range) to 1 (uniform coverage by an infinite number of points). A coverage of 0.9 indicates that the model’s variables, on average, have a maximum relative gap of 10%.

Gapsplit achieves better overall coverage with fewer samples than HR family algorithms. We generated samples from three genome-scale metabolic models using Gapsplit ACHR, and CHRR (Figure 1B). Both ACHR and CHRR plateaued within a few hundred samples at a coverage of 0.2, meaning each variable, on average, had an unsampled gap that covered 80% of the feasible space. Gapsplit samples quickly covered the solution space, reaching a coverage of 0.8 within 500 samples and plateauing with coverage over 0.9 after 2000 samples. The models were sampled as published with default bounds corresponding to glucose minimal media. We hypothesized that the fixed bounds prevented the HR samples from covering a larger fraction of the sample space. Indeed, opening all bounds to arbitrarily large values improved the coverage of the ACHR and CHRR samplers (Figure 1C). Gapsplit generated the best coverage for all models, but the HR samples also achieved coverage of at least 0.8. Thus Gapsplit gives better coverage of metabolic models especially when some of the variable bounds are fixed.

For models with fixed bounds, Gapsplit is slower than CHRR and faster than ACHR on a per sample basis (Figure 1D). However, Gapsplit is more efficient than either HR algorithm in the time required to reach a specific level of coverage since it yields better coverage per sample. Gapsplit is slower than either HR sampler for models with arbitrarily open bounds. However, we note that such models are not physiologically realistic since flux balance analysis requires at least one fixed constraint to limit nutrient uptake (Orth et al., 2010).

An advantage of Gapsplit is sampling non-convex models including those with discrete variables. We tested Gapsplit on two non-convex models of the yeast *Saccharomyces cerevisiae*: a metabolic model (Duarte et al., 2004) with gene-protein-reaction rules encoded as Boolean constraints (Shlomi et al., 2007; Jensen et al., 2011) and a combined metabolic/regulatory model (Herrgård et al., 2006). Adding hundreds of binary variables to the models did not affect the coverage during sampling (Figure 1E).

GapsplitŚ performance is tuned by changing only a single parameter: the number of secondary variables to target at each iteration. Changing this parameter (expressed as a fraction of the model’s total number of variables) can affect Gapsplit’s performance (Supplementary Figure S1). However, a single value (5%) was chosen as the default setting and worked well for all the experiments in this study. We do not expect users will need to tune this parameter for other models although they can change the parameter if needed.

## Conclusions

Gapsplit is a new class of random sampler for constraint-based models. It samples convex and non-convex models and outperforms HR family samplers on models with fixed bounds. Gapsplit is available in Matlab and Python and is compatible with models from the COBRA Toolbox (Heirendt et al., 2019), TIGER (Jensen et al., 2011), and cobrapy (Ebrahim et al., 2013). We believe Gapsplit opens new possibilities for exploring non-convex models including models with transcriptional regulation. Using Gapsplit, researchers can develop mixed-integer algorithms that incorporate random sampling.

## Acknowledgements

This work was supported by the National Institutes of Health grant EB027396. The authors declare no financial or commercial conflict of interest.

