## Supplementary Material for "gapsplit: Efficient random sampling for non-convex constraint-based models"

### The GAPSPLIT algorithm

GAPSPLIT begins with a COBRA model with $n$ variables $x_i, \ldots, x_n$. The model can be either convex (LP) or non-convex (MILP). GAPSPLIT calculates random samples by attempting to uniformly divide the solution space.

1. We use flux variability analysis (FVA) to find the feasible region for each variable: $x_i \in [\min(x_i), \max(x_i)]$. We define the $\text{range}(x_i) = \max(x_i) - \min(x_i)$. If the $\text{range}(x_i) < 1 \times 10^{-5}$, we consider the variable fixed and exclude it from targeting. We are left with $k \leq n$ non-fixed variables, which we will re-order in the model to be $x_1, \ldots, x_k$. In subsequent steps, we use the term "variables" to mean non-fixed variables that can be targeted by GAPSPLIT.

2. To find a new sample we select a single variable as the primary target. GAPSPLIT has three methods for choosing the primary target:

   - **Sequential** targeting chooses the variables in order (1 - $k$). After $k$ samples the algorithm cycles back to variable 1.
   - **Random** targeting chooses a variable at random for each sample.
   - **Max gap** targeting always selects the variable with the largest relative gap. If two or more variables have the same relative gap, one is chosen randomly from this subset.

   After the primary target is selected, we find its max gap. The new sample will split this gap evenly. If the max gap for variable $x_i$ spans [1.2, 2.4], then we set the lower and upper bounds for $x_i$ to be mean(1.2, 2.4) = 1.8. To avoid numerical issues, the actual bounds are set at ±0.1% of the targeted value.

3. GAPSPLIT guarantees the primary target's gap will be split evenly. GAPSPLIT also attempts to split a fraction of the other variables called secondary targets. By default GAPSPLIT selects 5% of the variables as secondary targets (Figure S1). If coverage is increasing slowly, the user may want to increase this fraction. If samples are taking too long to generate, the secondary fraction should be decreased. Assume the set $S$ contains the indices of the secondary targets. GAPSPLIT adds a quadratic objective to the model:

   $$\min_{x} \sum_{i \in S} \omega_i \left(x_i - \text{target}(x_i)\right)^2$$

   where $\text{target}(x_i)$ is the target point in the middle of the max gap of $x_i$ and $\omega_i = 1/\left(\text{range}(x_i)\right)^2$ is a regularization parameter that forces GAPSPLIT to weigh each variable equally regardless of its scaling.

4. The above QP (or MIQP) problem is solved to find a new sample point. The bounds of the primary target are reset and the max gaps for all variables are updated with the new sample.

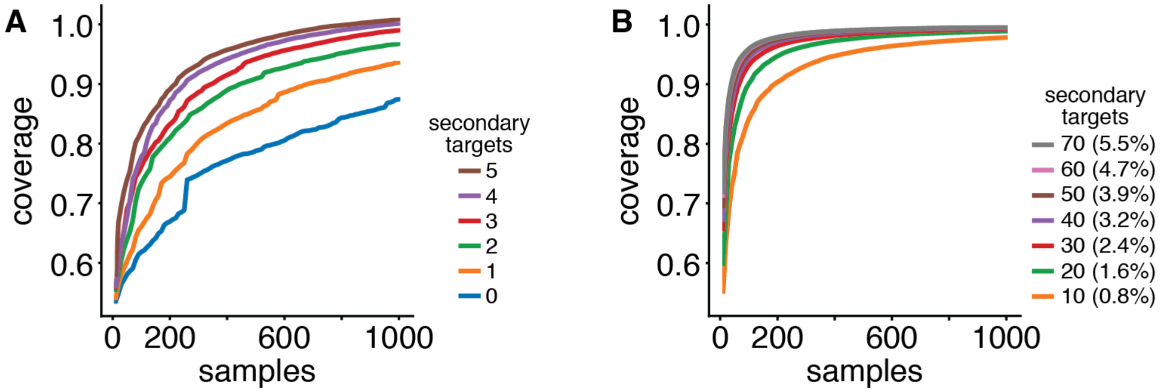


**Figure S1.** **A.** Increasing the number of secondary targets improves the coverage when sampling the yeast meatbolic model iND750. All simulations used sequential primary targeting and randomly selected secondary targets. **B.** The improvement in coverage levels off when the number of secondary targets exceeds 5% of the variables in the model. (The iND750 model has 1266 variables.) By default GAPSPLIT uses 5% of the model's variables as secondary targets.
